# Calcium Dependent Protein Kinase 3 is not essential for the asexual replication of *Plasmodium falciparum*

**DOI:** 10.64898/2025.12.04.692444

**Authors:** Himashree Choudhury, Utkarsh Gangwar, Nishtha Varshney, Aashukti Maheshwari, Subham Chintan Tripathy, Abhisheka Bansal

## Abstract

Calcium dependent protein kinases are essential at various stages of malaria parasite development within the vertebrate host and the mosquitoes. In the rodent malaria parasite, *Plasmodium berghei*, CDPK3 is essential for the sexual stage development of the parasite within mosquitoes. However, the function of CDPK3 in the most lethal human malaria parasite, *P. falciparum* remains unexplored. Moreover, the calcium dependent kinase activity of CDPK3 has not been experimentally tested. With the aim to biochemically characterize CDPK3 and perform its functional evaluation in the asexual replication of *P. falciparum*, here we have expressed the full-length CDPK3 in *E. coli* and demonstrated that it is a *bona fide* calcium dependent kinase using a luminescence-based assay and a semi-synthetic epitope approach. Complete knock-out of *cdpk3* using CRISPR/Cas9 did not show any defect in the asexual replication of the parasite and the knockout parasites look morphologically similar to the wild type. Our study suggests that *cdpk3* is not essential for the asexual replication of the parasite under *in vitro* culture conditions. Perhaps CDPK3 is required for the sexual development of *P. falciparum* like its ortholog in the rodent malaria parasite. CDPK3 may be a good target for the development of transmission blocking drugs.

## INTRODUCTION

Calcium ions (Ca²⁺) are universally used as secondary messengers to regulate a wide range of physiological and cellular processes in eukaryotes (1, 2). In *Plasmodium* species, Ca²⁺ modulates multiple signaling cascades critical for the development of malaria parasite at various stages in the vertebrate host and the mosquito (3). An important mechanism of transducing the intracellular calcium increase into cellular processes is mediated by the members of the calcium-dependent protein kinase (CDPKs) family. Upon binding of calcium ions, CDPKs are activated by overcoming the intramolecular inhibitory constraints imposed by a C-terminal junction domain. CDPKs have a multi-domain structure consisting of an N-terminal kinase domain followed by a calmodulin-like domain separated by a small junction domain. The junction domain acts as the autoregulatory domain that connects both the kinase domain and the calmodulin-like domain. The C-terminal calmodulin-like domain comprises four EF hand motifs and is important for the binding of calcium ions (4–6). *Plasmodium falciparum*, the most virulent malaria parasite, consists of seven members of CDPKs, i.e., CDPK (1–7) (7). Distinct CDPKs exhibit both unique and shared functions across different stages of the parasite’s life cycle, determined by their expression patterns, post-translational modifications, and variations in calcium-dependent substrate specificity and sensitivity. PfCDPK1 phosphorylates components of the motor complex and is critical for the red blood cell invasion (8–10). CDPK1 is also involved in schizont development and egress of merozoites from infected RBCs (11, 12). Apart from its essentiality for the blood stage development of the parasite, CDPK1 is also essential for the parasite transmission to mosquitoes. CDPK1 is critical for the formation of the male and female gametes (13). The second member of the CDPK family, CDPK2 is critical for the male gametocyte exflagellation and mosquito infection (14). CDPK2 knock-out parasites seems to form normal female gametes (14)unlike CDPK1 knock-out parasites that remain trapped in the host red blood cells (14). PfCDPK4 is critical for the sexual stage development of the parasite as inhibition of the kinase activity blocks the exflagellation process and transmission of the parasite to the mosquito vector (15–17). A phosphoproteomics study with the *cdpk4* knock-out parasites identified CDPK4 substrates in processes related to cell motility, DNA replication and translation (18). PfCDPK5 is essential for the egress of merozoites from mature schizonts. Conditional knock-down of CDPK5 using a destabilization domain arrested merozoites in the schizonts (19, 20). Interestingly, the trapped merozoites collected through mechanical rupture were capable of invading fresh RBCs suggesting that CDPK5 may not be required for the invasion of RBCs. PfCDPK7 is critical for the transition of ring to trophozoite during the blood stage of the parasite (21). Since CDPKs play essential roles in signalling pathways at various stages of the parasite life cycle and are absent in the human host (22), they are considered attractive drug targets for malaria treatment and prevention.

CDPK3, represents the third isoform and has been associated with various cellular functions in the rodent malaria parasite, *P. berghei*. PbCDPK3 is transcribed in the ookinete stage (23–25). Functional disruption of PbCDPK3 blocks the ookinete motility and subsequent invasion of the mosquito midgut. CDPK3 is essential for ookinetes to traverse the interface between the blood bolus and the midgut epithelial layer, enabling successful epithelial cell invasion (24). Ookinetes of *P. berghei* lacking functional CDPK3 exhibit a marked impairment in productive gliding motility, leading to a substantial decrease in their ability to transmit malaria to the mosquito vector (26). Although, the function of CDPK3 in rodent malaria parasite is known, however, no study has been conducted to investigate the role of CDPK3 (PF3D7_0310100) in the *P. falciparum*. Exploring CDPK3 function in the parasite biology may therefore provide better insight into the progression and pathogenesis of malaria. Moreover, molecular insights into the regulatory mechanisms of CDPK3 activation may open new avenues for designing drugs and inhibitory molecules to block its activity and therefore control parasite growth within RBCs.

In this study, we have expressed and purified full-length recombinant PfCDPK3 using affinity chromatography and verified its calcium-dependent kinase activity. We have successfully disrupted the endogenous *pfcdpk3* gene using clustered regularly interspaced short palindromic repeat (CRISPR)/CRISPR-associated protein 9 (Cas9) gene editing technique. *Pfcdpk3* knockout parasites exhibit normal asexual growth similar to that of wild-type (WT) parasites. Moreover, the morphology of the knockout parasite is also comparable to the WT parasite, suggesting that PfCDPK3 is likely non-essential for the asexual life cycle of the parasite but may have roles in other stages of the parasite’s life cycle.

## RESULTS

### PfCDPK3 is a *bona fide* calcium dependent protein kinase

CDPK3 conforms to the canonical domain architecture of the CDPK family containing an N-terminal variable domain followed by a kinase domain. At the C-terminus, a calmodulin-like domain (CamLD) containing four EF-hands is present. A junction domain separates the kinase domain from the CamLD (Fig.1a). The boundaries of the individual domains were found through pfam [http://pfam.xfam.org/]. To test the kinase activity of CDPK3 in the presence and absence of calcium ions, we expressed the full-length CDPK3 recombinant protein in *E. coli* with an N-terminal glutathione S-transferase (GST) tag. The CDPK3 recombinant protein was purified through affinity chromatography using glutathione sepharose resin. The purified protein conforms to the desired molecular weight of ∼ 91.6 kDa on the SDS-PAGE (Fig.1b). A difference in mobility is seen under non-reducing and reducing conditions on the SDS-PAGE (Fig. 1b). The recombinant protein was detected with the anti-CDPK3 antibodies generated in-house using CDPK3 CamLD as immunogen (Fig. S1). Under non-reducing condition, higher molecular weight bands were detected with the anti-CDPK3 antibody suggesting presence of CDPK3 multimeric forms (Fig. 1b). In addition to the desired full-length CDPK3 protein, several smaller fragments were detected both in the SDS-PAGE and the Western blot with anti-CDPK3 antibodies suggesting degradation of the protein during purification.

**Figure 1.**
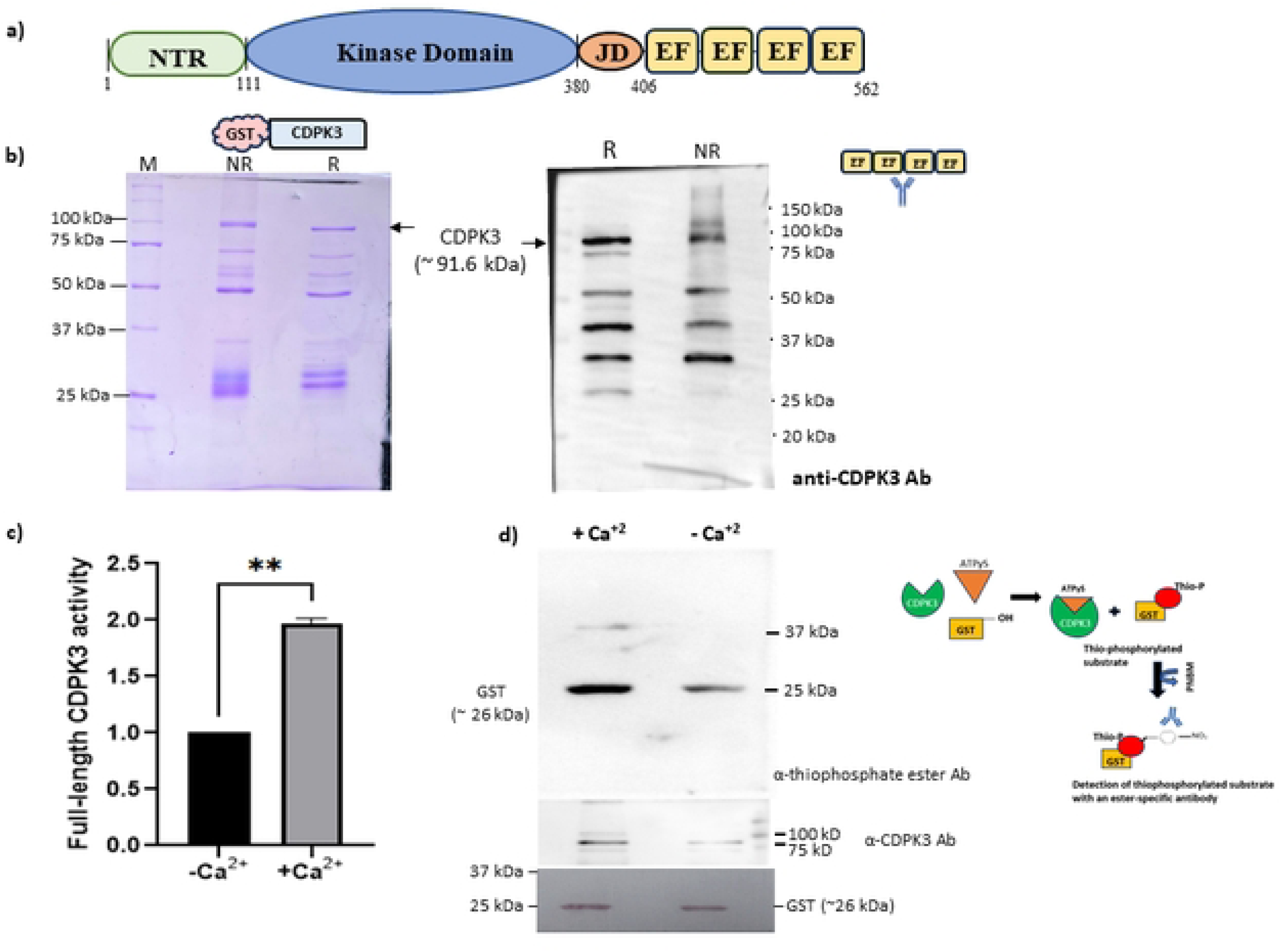
CDPK3 is a *bona fide* calcium dependent protein kinase. **a)** The schematic shows the domain architecture of the full-length CDPK3. CDPK3 contains an N-terminal domain (NTR) followed by a kinase domain. A junction domain (JD) separates the kinase domain from the C-terminal calmodulin-like domain containing calcium binding motifs called EF-hands. The boundaries of each domain are indicated [http://pfam.xfam.org/]. **b)** SDS-PAGE and Western blot analysis of full-length CDPK3 tagged with an N-terminal glutathione S-transferase (GST). Coomassie R250 stained SDS-PAGE shows the purified recombinant CDPK3 under reducing (R) and non-reducing (NR) conditions. M- unstained protein ladder. (right) Western blot of the recombinant protein under R and NR conditions with anti-CDPK3 antibody shows the detection of the full-length CDPK3 along with few truncated fragments. **c)** Recombinant CDPK3 shows calcium dependent increase in kinase activity. *In vitro* kinase assay with recombinant CDPK3 was set in absence (-Ca^2+^) and presence (+Ca^2+^) of calcium chloride and the enzyme activity was measured using the luminescence-based ADP-Glo kinase assay kit. The activity of the enzyme is plotted on the Y-axis in the -Ca^2+^ and +Ca^2+^ conditions. The increase in the kinase activity of full-length CDPK3 is statistically significant (**p=0.003, unpaired t test with Welch’s correction, n=3). The error bars show the standard error of the mean. The graph is plotted with GraphPad Prism 10.5.0. **d)** A Western blot format shows calcium dependent increase in the transphosphorylation activity of CDPK3. A semi-synthetic epitope tagging approach (shown as a schematic on the right) was used to test the kinase activity of CDPK3 in the presence (+Ca^2+^) and absence (-Ca^2+^) of calcium using Glutathione S-transferase (GST) as an exogenous substrate. Thiophosphorylated reaction products were detected using anti-thiophosphate ester specific antibody. The reaction mixtures were also probed with anti-CDPK3 antibody as loading controls under the two conditions. Ponceau stained PVDF membrane after transfer of proteins shows loading of GST under the two conditions.

The CDPK3 recombinant protein was used to set up *in vitro* kinase assay using Myelin Basic Protein (MBP) as an exogenous substrate in the presence (+ 5 mM CaCl_2_) and absence (+ 2.5 mM EGTA) of calcium ions. Kinase activity of the recombinant protein was detected using ADP-Glo assay (27). Luminescence readout, reflecting the conversion of ATP to ADP by the recombinant CDPK3, was higher in the presence of calcium than in its absence (Fig. 1c). The increase in the kinase activity of CDPK3 (1.965 ± 0.053) in the presence of calcium was statistically significant (p=0.003, unpaired t test with Welch’s correction, n=3) (Fig. 1c) suggesting that the recombinant CDPK3 is a calcium-dependent protein kinase.

To further confirm the calcium dependent activation of CDPK3, we used a semi-synthetic epitope tagging approach (28). The kinase assay reaction was set-up with the recombinant CDPK3 in the absence and presence of calcium chloride using GST as an exogenous substrate. ATPγS, a ATP analog, containing terminal thiophosphate group was used as the source of the transferable phosphate. The thiophosphorylated residues were alkylated using p-nitrobenzyl mesylate (PNBM) and detected with a specific anti-thiophosphate ester antibody. Transphosphorylation of GST was substantially increased in the presence of calcium compared to without calcium condition (Fig. 1d). An additional band around 37 to 50 kDa was detected in the presence of calcium (Fig. 1d) that may be the truncated fragment of the full-length CDPK3 also visible on the SDS-PAGE (Fig. 1a). We did not detect the autophosphorylated CDPK3 suggesting that either CDPK3 is not autophosphorylated or the autophosphorylated CDPK3 is beyond the detection limit of the assay format. Taken together, our data shows that the recombinant CDPK3 is a *bona fide* calcium dependent protein kinase.

### *cdpk3* can be completely knocked-out from the asexual blood stages

An initial study showed that the transcript of CDPK3 is present only in the sexual stages of *P. falciparum* (29). However, transcriptomics data in PlasmoDB shows CDPK3 expression in the blood stages [PlasmoDB.org,(30–32)]. In order to elucidate the function and test essentiality of *cdpk3* for the blood stage replication of the parasite, we used the CRISPR/Cas9 gene editing to completely disrupt the *cdpk3* locus. A single plasmid, *pL6-Cas9-cdpk3ko*, encoding Cas9 endonuclease, *cdpk3* specific sgRNA, the repair sequence containing the 5’ and the 3’ homologous sequences was generated (Fig. 2a). Drug resistant parasites obtained after the transfection were confirmed by a diagnostic PCR for the desired modification. Genetic material prepared from the WT and three independent clones of *cdpk3ko* parasites (*Pf:cdpk3^ko^*): A4, C5, and D5, was amplified using different primer sets. Primers, F1 and R1, corresponding to the 5’ and 3’ utr regions of *cdpk3*, respectively were designed that were not part of the repair DNA template. A reverse primer, PL6_16 corresponding to the hDHFR sequence was used along with F1 for the PCR. An amplicon of 660 bp was detected in all the clones of the *Pf:cdpk3^ko^*parasites with the primer set F1/PL6_16 while no amplification was observed in the WT parasite (Fig. 2b). Amplification of the entire region across the *cdpk3* homologous sequences with F1/R1 resulted in amplicons of different sizes in the WT (2897 bp) and the *Pf:cdpk3^ko^* parasites (3110 bp). The WT contamination in the *Pf:cdpk3^ko^* parasites was tested with the primer set F2/R1. F2 corresponds to the region deleted in the *Pf:cdpk3^ko^* parasites. An expected amplicon of 813 bp was detected in the WT parasites while there was no specific amplification in the *Pf:cdpk3^ko^* clones (Fig. 2b).

**Figure 2.**
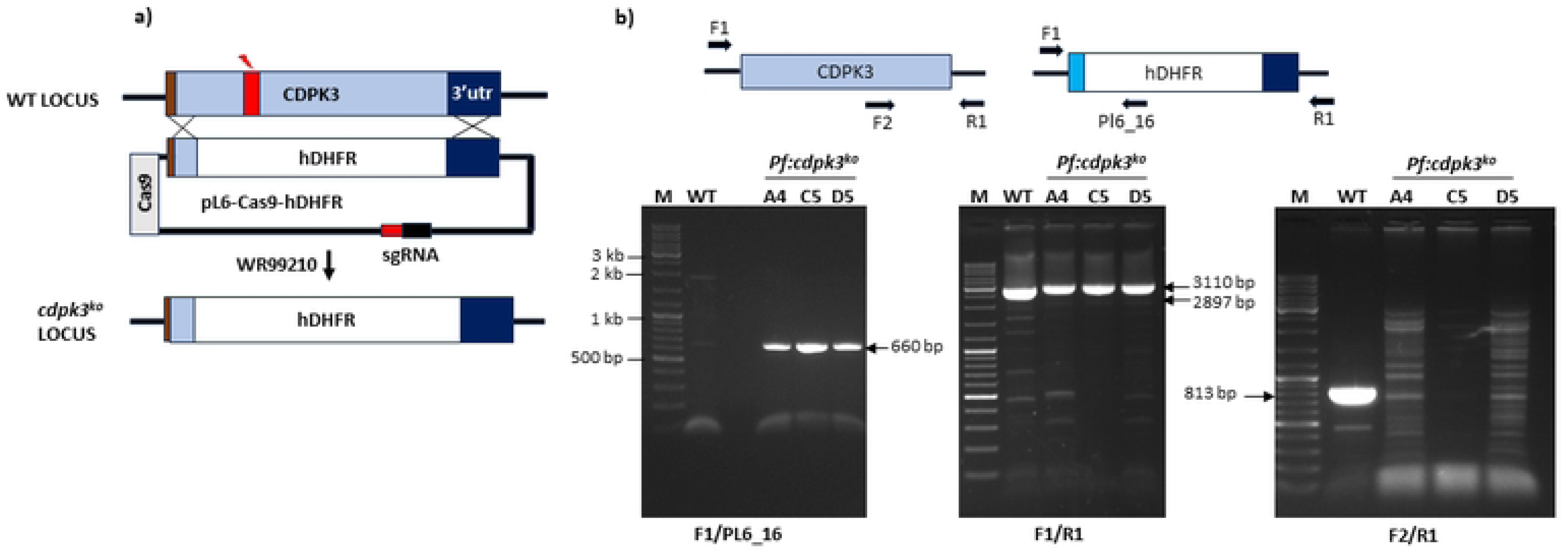
Verification of *cdpk3* knock-out in the asexual blood stages. **a)** Scheme representing the strategy to knock-out endogenous *cdpk3* in the parasite. A region in the endogenous *cdpk3* locus is selected for Cas9 endonuclease cleavage (red box). The CRISPR/Cas9 plasmid, pL6-Cas9-hDHFR, expresses sgRNA, Cas9 endonuclease, and the hDHFR drug-resistance cassette. The hDHFR cassette is flanked by the 5’ and the 3’ homologous regions corresponding to the wild type *cdpk3* sequence. The sgRNA consists of a 20-nucleotide guide sequence (red box) and the tracrRNA. Parasites transfected with pL6-Cas9-hDHFR plasmid are selected with WR99210. After the desired editing, the DNA sequence following the 5’ homologous sequence is replaced by the hDHFR cassette resulting in complete knockout of *cdpk3* (*cdpk3^ko^*). **b)** PCR verification of the *cdpk3* knockout clones. Three independent clones of *cdpk3* knock-out, *Pf:cdpk3^ko^* A4, C5, and D5 were used to verify desired editing of the endogenous locus. Amplification with the primer set, F1/PL6_16, show amplicons of expected size (660 bp) only in the *Pf:cdpk3^ko^* parasites. Amplification with the primer set, F1/R1 show difference in the size of the amplicons in the WT (2897 bp) and the *Pf:cdpk3^ko^* (3110 bp) parasites. Absence of the WT contamination in the *Pf:cdpk3^ko^* clones was verified with the wild type specific primer set, F2/R1. A specific amplicon of expected size (813 bp) is observed only in the WT parasites and not in the *Pf:cdpk3^ko^* clones. M- molecular weight marker.

Clone C5 of the *Pf:cdpk3^ko^* parasites was further verified for the complete knock-out of *cdpk3* using additional sets of primers. Three sets of primers: F2/R2, F4/R2, and F3/R2, specific only for the WT parasites showed amplicons of the expected sizes of 435 bp, 1351 bp and 1939 bp, respectively only in the WT parasites while no amplification was observed in the *Pf:cdpk3^ko^* C5 (Fig. S2). DNA template from both the parasites were used for the targeted amplification of *cdpk1* and *cdpk5* genes using gene specific primers. Amplicons of the expected sizes of 2225 bp and 1700 bp were obtained for *cdpk1* and *cdpk5*, respectively in both the parasites (Fig. S2). These results confirm successful disruption of *cdpk3* locus in the knock-out parasites.

### *Pf:cdpk3^ko^* parasites show no perceptible growth defect under *in vitro* culture conditions

Our results show that the mutant parasites with disrupted *cdpk3* locus are viable and can grow under *in vitro* conditions. To check whether the disruption of *cdpk3* cause any growth defects in the *Pf:cdpk3^ko^* parasites, we compared the asexual replication of the knockout parasites with the WT for 5 consecutive days. Highly synchronized trophozoite stage parasites were seeded at 0.5 % parasitemia and the growth of the parasites was monitored every day through SYBR Green I assay and Giemsa stained smears. The growth of the *Pf:cdpk3^ko^* parasites was comparable to the WT (Fig. 3a & Fig. S3). The Giemsa stained smears were used to check the morphology of different asexual stages of the *Pf:cdpk3^ko^* and the WT parasites. No perceptible difference was seen in the morphology of the *Pf:cdpk3^ko^*parasites compared to the control in the 1^st^ and the 2^nd^ replication cycle (Fig. 3b). These results suggest that CDPK3 is redundant for the asexual proliferation of the parasite under *in vitro* conditions.

**Figure 3.**
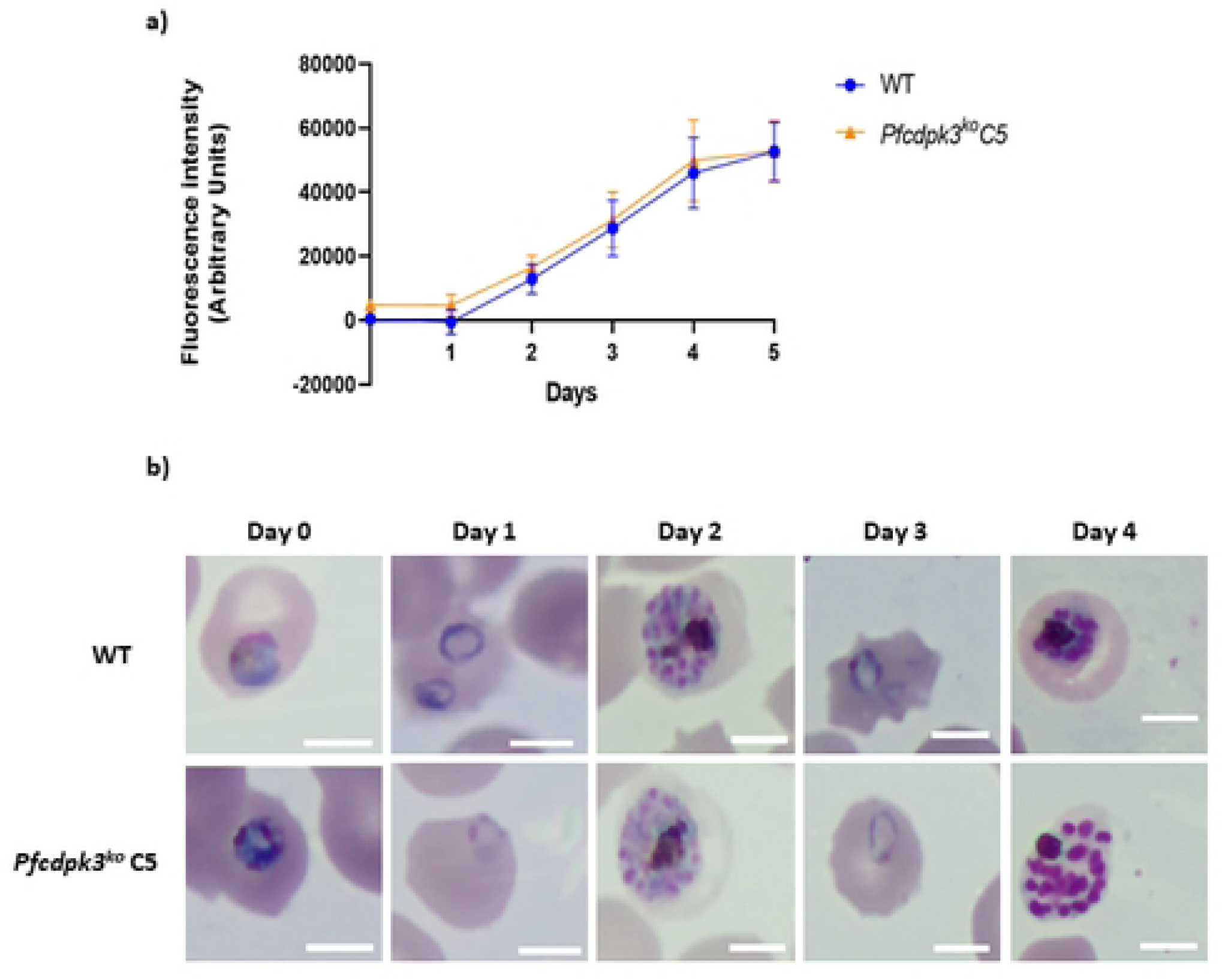
*Pf:cdpk3^ko^* parasites show no growth defect under *in vitro* culture conditions. **a)** A SYBR Green I assay was used to test the asexual proliferation of the *Pf:cdpk3^ko^* C5 clone along with the parental WT parasites. Tightly synchronized parasites were seeded at equal parasitemia and the growth was monitored by sampling the parasites every day for 5 days. Fluorescence Intensity, expressed as arbitrary units, on the Y-axis is plotted against number of days on the X-axis. Data from three independent experiments performed in triplicate is used to plot the graph through GraphPad Prism. The error bars represent the standard deviation of the mean intensity. The asexual proliferation of another independent clone of *Pf:cdpk3^ko^* i.e., *Pf:cdpk3^ko^* D5 was also tested through SYBR Green I assay along with the WT (Fig. S3). **b)** The *Pf:cdpk3^ko^* parasites show no morphological defects in the asexual blood stages. Giemsa stained pictures of different stages of the *Pf:cdpk3^ko^* C5 and WT parasites were acquired for Day 0 through Day 4 with the 100X objective in a bright field microscope. Representative pictures of each parasite are shown for the corresponding days. The scale bars correspond to 5 μm.

## MATERIAL AND METHODS

### Molecular cloning and expression of full length CDPK3 (FLCDPK3)

*P. falciparum* (PF3D7_0310100) CDPK3 gene full-length sequence was retrieved from PlasmoDB [Plasmodb.org]. The complete CDS of *cdpk3* gene was amplified with PrimeSTAR Max DNA polymerase (TAKARA Bio) using the primer pair: CK3FpgexFwd / CK3FpgexRev. The complementary DNA (cDNA), prepared from asynchronous parasite culture, was used as a template since introns are present in the *cdpk3* gene sequence. The PCR conditions used for the amplification are: 98 °C/2 min, 36 cycles of (98 °C/10 sec, 52 °C/20 sec, and 62 °C/45 sec), and final extension at 62 °C/10 min. The amplified product was digested with the restriction enzymes BamHI/NotI along with the target plasmid, pGEX4t1. The construct was designed to express full length CDPK3 with an N-terminal Glutathione S-Transferase (GST) Tag. The plasmid and the insert were ligated with T4 DNA ligase 16°C/16 h and transformed into *E. coli* DH5α competent cells. Positive clones containing the plasmid with the desired insert were confirmed by double-digestion with BamHI/NotI and DNA sequencing.

To express CDPK3 recombinant protein, *E. coli* BL21 (DE3)pLysS strain was transformed with the sequence verified plasmid and selected with appropriate combination of antibiotics. A single clone was used to set up the primary culture followed by the secondary culture. The secondary culture was grown to 0.7 OD_600_ in a shake flask at 37 °C with ampicillin & chloramphenicol as selection markers. The recombinant protein expression was induced with 1 mM IsoPropylThioGalactoside (IPTG) and grown further at 30°C/5 h. The *E. coli* cells were harvested at 6000 g for 5 min at 4 °C post-induction and lysed in a buffer with composition: 50 mM Tris (pH 7.0) and 100 mM NaCl, 1 mM PMSF (phenylmethylsulfonyl fluoride), 20 mM MgCl_2_, 2 µM EDTA, 0.2 % Triton X 100, 10 % glycerol & 0.1 mg/mL lysozyme, followed by sonication for 3 min (10 % amplitude with Pulse ON: 8 sec, Pulse Off: 10 sec). The lysate was then centrifuged at 9,500 rpm for 1 h at 4 °C. The supernatant containing recombinant CDPK3 protein was incubated with Glutathione Sepharose resin pre-equilibrated with the buffer containing 50 mM Tris, pH 7.0 & 100 mM NaCl overnight at 4 °C. The resin was allowed to settle by centrifugation at 500 g for 5 min at 4 °C followed by extensive washing (7 times) using wash buffer (50 mM Tris, pH 7.0 and 100 mM NaCl) at 500 g for 3 min to remove the unbound proteins. The bound CDPK3 protein was eluted from the Glutathione Sepharose resin with 10 mM & 20 mM reduced glutathione in the wash buffer. The fractions were run on 10 % SDS-PAGE. The pure fractions were stored at −80 °C for downstream experiments.

### *In vitro* ADP-Glo Kinase assay with the recombinant CDPK3 protein

To test the calcium dependent kinase activity of recombinant CDPK3 protein, we used an ADP-Glo Kinase assay platform. 300 ng of recombinant CDPK3 protein was incubated with the kinase reaction buffer containing 40 mM Tris-Cl, pH-7.5, 20 mM MgCl_2_, 500 mM PhIC (Phosphatase Inhibitor Cocktail), 100 µM ATP (Ultra-Pure) and 10 ng of Myeline Basic Protein, MBP used as an exogenous substrate for 50 µl of reaction. The kinase reactions were set either in the presence [+ 5 mM CaCl_2_] or absence [+ 2.5 µM EGTA) of calcium and incubated at 30 °C for 1 h in a stirred water bath. After the completion of the kinase reaction, the ADP-Glo™ Reagent & the Kinase Detection Reagent were added in equal and double the volume of the kinase reaction, respectively. The final ratio of the Kinase Reaction: ADP- Glo^TM^ Reagent: Kinase Detection Reagent is 1:1:2. The luminescence was recorded, and the standard curve was plotted to estimate the kinase activity of the recombinant CDPK3 protein.

A standard curve was also generated with each experiment to precisely calculate the ADP to ATP ratio in each experimental condition. First, a kinase reaction buffer is prepared containing 40 mM Tris-Cl, pH-7.5, 20 mM MgCl_2_, 0.1 mg/mL BSA. Different % of ADP: ATP were used to generate a standard curve for percent ATP to ADP conversion using Ultra-Pure ATP (10 mM) & ADP (10 mM). Using serial dilution, ATP & ADP final concentration was made 100 µM. To 5 µl of the kinase reaction mixture, 5 µl of ADP-Glo™ Reagent was added and incubated for 30 min at RT. Following this, 10 µl of Kinase Detection Reagent was added and incubated further for 40 min at RT. The luminescence read-out was taken on FLUOstar Omega Filter-based multi-mode microplate reader.

### Semi-synthetic epitope approach to detect kinase activity of CDPK3

The activity of recombinant CDPK3 was assessed through an *in vitro* kinase assay employing a semisynthetic epitope tagging method, as outlined by Allen et al (28). Approximately 200 ng recombinant protein was incubated in a buffer composed of the following components: 50 mM Tris, 50 mM MgCl_2_, 1x phosphate inhibitor cocktail (Roche PhosSTOP™) with an exogenous substrate, Glutathione S-transferase (GST). The buffer was prepared under conditions either requiring the presence of calcium (5 mM CaCl₂) or its absence, achieved by the addition of 2.5 mM EGTA. Importantly, 100 μM ATPγS was used as the source of the transferable phosphate group. The reaction mixture was incubated at 30°C for 2 h, followed by terminating the reaction by the addition of 5 mM EGTA. A 5 mM concentration of p-nitrobenzyl mesylate (PNBM) was added to the reaction mixture and incubated at 20°C for 2 h to allow protein alkylation. The reaction was stopped by 1x SDS sample buffer. Thiophosphorylated substrates were detected through Western blot using anti-thiophosphate ester antibody (Ab92570, abcam).

### Molecular cloning and expression of CDPK3 calmodulin-like domain

The four EF-hands containing calmodulin-like domain of *cdpk3* (1252-1686 nucleotides) was amplified using cDNA with the primer set: Ck3efpET42F/Ck3efpET42R. The same amplification conditions were employed as described above for the full-length construct of *cdpk3* except for the cyclic amplification at 62 °C/30 sec. The amplified product was digested with NdeI/XhoI along with the plasmid pET42a (+). The plasmid and the insert were ligated with T4 DNA ligase 16 °C/16 h and transformed into XL10- Gold competent cells. Positive clones containing the plasmid with the desired insert were confirmed by were double-digestion with NdeI/XhoI and DNA sequencing.

To express CDPK3-CamLD, the *E. coli* ArcticExpress (DE3) competent cells were transformed with the sequence verified clone. An isolated clone from the transformed *E. coli* cells was inoculated in 5 mL Luria Bertani broth with appropriate antibiotics and incubated O/N at 37 °C with constant shaking at 220 rpm. A secondary culture was set from the primary culture and incubated at 30 °C, shaking at 220 rpm for around 3 h till 0.7 OD_600_ is reached. The expression of CDPK3-CamLD was induced with 1 mM IPTG and the flask was further incubated at 12 °C/24 h. The recombinant CDPK3-CamLD contains 8x HIS tag at the C-terminus with an estimated molecular weight of ∼ 18 kDa. CDPK3-CamLD was purified using the methodology as described for the CDPK1-CamLD (8).

### Antibody generation against CDPK3

To generate CDPK3-specific antibodies, we used CDPK3-CamLD protein as the antigen of choice as the primary amino acid sequence of calmodulin-like domain is very specific in different CDPKs. Antibodies against CDPK3 were generated in-house in Balb/c mice as described previously (14). Pre-bleed was collected from the mice 2 days before the first immunization. Priming was done on Day 0 with 50 μg of CDPK3-CamLD in Complete Freund’s Adjuvant. The 1^st^ and the 2^nd^ booster were given with 25 μg of CDPK3-CamLD formulated in Incomplete Freund’s Adjuvant after 21 and 42 days of the priming, respectively. After 1 week of the 2^nd^ boost, terminal bleed was collected. Serum containing anti-CDPK3-specific antibodies was isolated from whole blood by allowing the blood to clot for 30-40 minutes at room temperature, followed by centrifugation at 2000 rpm for 10 minutes at 4 °C. The collected antiserum was stored in small aliquots at −80 °C for later usage.

The Ethical clearance was obtained from the Institutional Animal Ethics Committee (IAEC code # 12/2024) before proceeding with the animal use. The procedures were conducted in the Central Laboratory Animal Resources (CLAR), Jawaharlal Nehru University following the guidelines of the Committee for Control and Supervision of Experiments on Animals (CCSEA).

### In vitro culture of P. falciparum

*P. falciparum* (NF54 strain) was propagated at 2 % hematocrit in O+ human RBCs (Rotary Blood Bank, New Delhi, India). RPMI1640 medium containing L-glutamine was supplemented with 25 mM HEPES (Sigma-Aldrich), 50 µg/mL Hypoxanthine (Sigma-Aldrich), 0.5 % AlbuMAX II Lipid-Rich BSA (ThermoFisher Scientific), 25 mM sodium bicarbonate (Sigma-Aldrich) and 10 μg/mL gentamycin (ThermoFisher Scientific) as described earlier (33). Culture flasks containing the parasitized RBCs were flushed with sterile mixed gas of the following composition: 5 % O_2_, 5 % CO_2_, and 90 % N_2_ at 37 °C. Experiments requiring mature schizont stage parasites were obtained through two consecutive cycles of sorbitol.

### Generation of *cdpk3* knockout parasites using CRISPR/Cas9

To delete *cdpk3* from the endogenous locus, we employed CRISPR/Cas9 gene editing. An appropriate guide DNA sequence (ATGAAATGAAGACCAAAGCA) was cloned in the pL6-Cas9-hDHFR plasmid with the oligo pair Gcdpk3F/Gcdpk3R using In-Fusion kit (Takara Bio), following the manufacturer’s instructions. Subsequent to the cloning of the guide sequence, the 5’ and 3’ homology fragments were cloned. The 5’ homology fragment (405 bp) was amplified using the primer pair: 5HRCDPK3F/5HRCDPK3R and cloned within the SacII/AflII restriction enzyme sites. The 3’ homology fragment was amplified with the primer set: 3HRCDPK3F/3HRCDPK3R and cloned within EcoRI/NcoI restriction enzyme sites. All the cloned elements were verified with DNA sequencing. The complete construct, *pL6-Cas9-cdpk3ko*, containing all the essential elements for CRISPR mediated knockout of *cdpk3* was used to transfect the ring stage parasites using electroporation as described earlier (34, 35). Following transfection, the parasites were selected with 2.5 nM WR99210 till the appearance of the drug resistant parasites. After confirming the presence of the parasites with the desired editing by PCR, limiting dilution cloning was set up to obtain the clonal parasites. The clonal parasites were further confirmed for the desired editing with appropriate set of primers and the contamination of the wild type parasites was ruled out using primers specific for the *cdpk3* WT locus. Clone C5 of the CDPK3 knock-out parasite, *Pf:cdpk3^ko^* C5, was further verified for the desired editing of the *cdpk3* locus with additional sets of primers that specifically amplify only the WT locus and not the disrupted locus. The *Pf:cdpk3^ko^* C5 parasite was used for the asexual growth experiment and morphological analysis through bright field microscopy.

### *In vitro* malaria parasite growth assay and light microscopy

Wild-type and *Pf:cdpk3^ko^* clonal parasites (*Pf:cdpk3^ko^* C5 and *Pf:cdpk3^ko^* D5) were sorbitol-synchronised for two consecutive cycles, followed by seeding the parasites at 0.2% trophozoite stage (30-36 hr post invasion) in a 24-well plate. The parasites were allowed to grow under optimal conditions as mentioned in the “*In vitro* culture of *P. falciparum*”. The Giemsa slides were prepared for 5 days and assessed under the microscope to monitor the asexual growth progression for up to 5 days. Parasite samples were also drawn every 24 h for 5 days and stored in −80°C to further evaluate growth using SYBR Green I-based fluorescence assay. The frozen parasite samples were thawed and plated in a 96- well plate and lysed in a lysis buffer of the following composition: Tris (20 mM; pH 7.5), EDTA (5 mM), and saponin (0.008%; wt/vol). The plate was incubated at 37°C for half an hour. Later, the fluorescence readings were acquired at the excitation and emission wavelengths of 485 nm and 520 nm, respectively, in Fluostar Optima (BMG Labtech) plate reader with a gain of 1000. Experiments were conducted in triplicate, with two independent repeats for *Pf:cdpk3^ko^* D5 and three independent repeats for *Pf:cdpk3^ko^*C5. The growth curve was generated by plotting the number of days (x-axis) against the fluorescence (Y-axis) using GraphPad Prism 8.

Microscopic images of *Pf:cdpk3^ko^* C5 and the WT parasites were acquired on the Zeiss Primostar Trinocular Microscope with axiocam 208 color camera using 100X oil immersion objective.

## DISCUSSION

Autophosphorylation is used by some kinases as a regulatory mechanism to control activity (36, 37). CDPKs show autophosphorylation activity that may either increase (36) or decrease the substrate level phosphorylation (38). Autophosphorylation of the variable N-terminal domain has been demonstrated for other CDPKs that modulates the interaction with other proteins (39–41). We did not observe autophosphorylation of recombinant CDPK3 under the conditions employed in this study suggesting that unlike other members of the CDPK family, PfCDPK3 does not undergo autophosphorylation (8, 37, 42, 43). It may be possible that the autophosphorylation signals are too weak to be detected using the semi-synthetic epitope tag approach. The ATP analog, ATPγS, utilized to visualize the auto- and transphosphorylation activities of recombinant CDPK3 through the semi-synthetic approach is hydrolysed much more slowly than the normal ATP and may also be a reason for not observing the autophosphorylated CDPK3. Additionally, the calcium-dependent activity of CDPK3 is predictably lower compared to other members of the *Plasmodium* CDPK family since one of the EF-hands does not have the conserved residues required for calcium ion binding (5). It has been shown that the N-lobe of the CDPK3 CLD can bind only a single calcium ion unlike the C-lobe that already contains two calcium ions bound even in the inactive state (44). The activation of CDPK3 therefore would require much higher concentrations of calcium than other CDPKs of *Plasmodium*. The overall structural changes requiring activation of CDPK3 in the presence of calcium are initiated by the binding a single calcium ion on the N-lobe of CLD. This results in the movement of the complete CAD away from the kinase domain relieving the autoinhibitory constraints due to the junction domain (6). The activation mechanism of CDPKs is not dependent on the phosphorylation of activation loop due to the presence of conserved acidic residues (45).

The transcript of *cdpk3* was not detected in the asynchronous culture of *P. falciparum* (29). However, few other studies suggest that the transcript of *cdpk3* is present in the asexual blood stages (30–32). Furthermore, the inability to completely disrupt *cdpk3* led to the suggestion that the gene may be likely essential for the blood stages (46). In order to directly test the essentiality of *cdpk3* for the blood stage parasite, we used CRISPR/Cas9 tool to completely knock-out the gene. We did not see any perceptible difference in the morphology of the CDPK3 knock-out parasites during the asexual growth. Moreover, the knockout parasites grew as well as the WT suggesting that CDPK3 is perhaps redundant for the asexual stages. Complete disruption of PbCDPK3, an ortholog of PfCDPK3 did not show any defect during asexual growth (24, 26). The cdpk3ko parasites could not transmit to the mosquitoes due to defect in ookinete motility and traversal of midgut epithelium (24, 26). *P. falciparum* CDPK3 may be essential during the sexual stage development within the mosquito like *P. berghei*.

CDPK4 is not essential for the asexual replication of the parasite however, in the background of mutant PKG (T619Q), the *cdpk4* knock-out parasites show significant reduction in the asexual growth compared to the single mutant parasites (47). The two genes therefore show negative epistasis and the function of CDPK4 becomes important for the invasion of RBCs during the asexual replication in the mutant parasite background (47). Likewise, although CDPK3 is not essential for the asexual growth of the parasite however, it may also show negative interactions with other proteins that are important for the asexual replication. This possibility may be explored in future studies that may provide useful insights into the epistatic interactions of *cdpk3* with other kinases. At present, CDPK3 cannot be used as a target for blood stages however, it might be critical for the sexual stage development of *P. falciparum* in mosquitoes like PbCDPK3. The function of CDPK3 needs to be tested in the sexual development within mosquitoes.

---

**Figure S1. SDS-PAGE and Western blot analysis of calmodulin-like domain of CDPK3.** The C-terminal calmodulin-like domain of CDPK3 was expressed with a C-terminal 8XHIS tag. The purified recombinant CDPK3-CamLD was separated on a 12 % SDS-PAGE under non-reducing (NR) and reducing (R) conditions and stained with Coomassie-R250. The recombinant CDPK3-CamLD predominantly exists as a monomer (∼ 18 kDa) and dimer (∼ 36 kDa) under reducing and non-reducing conditions, respectively. Western blot with anti-HIS and anti-CDPK3 antibodies show the detection of the recombinant protein corresponding to the SDS-PAGE profile. M- unstained molecular weight marker.

**Figure S2. PCR verification of the *cdpk3* knockout clone 5, *Pf:cdpk3^ko^* C5.** The *Pf:cdpk3^ko^* C5 was verified with additional sets of primers to confirm complete disruption of the *cdpk3* locus. Amplification with the primer sets, F2/R2, F2/R1, F4/R2, and, F3/R2 show amplicons of expected sizes, 435 bp, 813 bp, 1351 bp, and 1939 bp respectively only in the WT parasite. (right) *Pf:cdpk3^ko^* C5 shows comparable amplification of the *Pfcdpk1* (2225 bp) and *Pfcdpk5* (1700 bp) loci as the WT. M- molecular weight ladder.

**Figure S3. *Pf:cdpk3^ko^* parasites show no growth defect under *in vitro* culture conditions.** SYBR Green I assay was used to test the asexual proliferation of 2 independent clones of *Pf:cdpk3^ko^* i.e., C5, and D5 along with the parental WT parasite. Tightly synchronized parasites were seeded at equal parasitemia and the growth was monitored by sampling the parasites every day for 5 days. Fluorescence Intensity, expressed as arbitrary units, on the Y-axis is plotted against number of days on the X-axis. Data from two independent experiments performed in triplicate is used to plot the graph through GraphPad Prism. The error bars represent the standard deviation of the mean intensity. The asexual proliferation of *Pf:cdpk3^ko^*C5 from 3 independent experiments performed in triplicate is shown separately as Fig. 3.

## ACKNOWLEDGEMENTS

Financial support from the University Grants Commission (grant # F.30-433/2018(BSR)) and Department of Biotechnology (grant # BT/PR38411/GET/119/311/2020) is acknowledged. Facilities/laboratories supported by DBT-Builder (BT/INF/22/SP45382/2022) is also acknowledged. The funding agency has no role in the preparation and decision to publish the work. We acknowledge the support received from the animal facility of JNU, Central Laboratory Animal Resources (CLAR) for the generation of antisera against CDPK3. We acknowledge Jacobus Pharmaceutical Company, Inc and Kristin D. Lane for WR99210. We thankfully acknowledge Ashis K. Nandi for allowing the usage of Fluostar Optima (BMG Labtech) plate reader to read the 96-well plates for SYBR Green-I assay and the ADP-Glo Kinase assay.

## AUTHORS CONTRIBUTIONS

**Conceptualization:** Abhisheka Bansal; **Methodology:** Himashree Choudhury, Utkarsh Gangwar, Nishtha Varshney, Aashukti Maheshwari, Subham Chintan Tripathy; **Formal analysis and investigation:** Himashree Choudhury, Utkarsh Gangwar, Nishtha Varshney, Aashukti Maheshwari,; **Writing - original draft preparation:** Abhisheka Bansal, Himashree Choudhury; **Writing - review and editing:** Abhisheka Bansal, Himashree Choudhury; **Funding acquisition:** Abhisheka Bansal; **Resources:** Himashree Choudhury, Utkarsh Gangwar, Nishtha Varshney, Aashukti Maheshwari, Subham Chintan Tripathy; **Supervision:** Abhisheka Bansal. All authors reviewed the manuscript.

## REFERENCES

1. Carafoli E, Krebs J. Why Calcium? How Calcium Became the Best Communicator. J Biol Chem. 2016;291(40):20849–57.

2. Cremer T, Neefjes J, Berlin I. The journey of Ca(2+) through the cell - pulsing through the network of ER membrane contact sites. J Cell Sci. 2020;133(24).

3. de Oliveira LS, Alborghetti MR, Carneiro RG, Bastos IMD, Amino R, Grellier P, et al. Calcium in the Backstage of Malaria Parasite Biology. Front Cell Infect Microbiol. 2021;11:708834.

4. Huang JF, Teyton L, Harper JF. Activation of a Ca(2+)-dependent protein kinase involves intramolecular binding of a calmodulin-like regulatory domain. Biochemistry. 1996;35(40):13222–30.

5. Wernimont AK, Amani M, Qiu W, Pizarro JC, Artz JD, Lin YH, et al. Structures of parasitic CDPK domains point to a common mechanism of activation. Proteins. 2011;79(3):803–20.

6. Wernimont AK, Artz JD, Finerty P, Jr., Lin YH, Amani M, Allali-Hassani A, et al. Structures of apicomplexan calcium-dependent protein kinases reveal mechanism of activation by calcium. Nat Struct Mol Biol. 2010;17(5):596–601.

7. Sharma M, Choudhury H, Roy R, Michaels SA, Ojo KK, Bansal A. CDPKs: The critical decoders of calcium signal at various stages of malaria parasite development. Comput Struct Biotechnol J. 2021;19:5092–107.

8. Bansal A, Singh S, More KR, Hans D, Nangalia K, Yogavel M, et al. Characterization of Plasmodium falciparum calcium-dependent protein kinase 1 (PfCDPK1) and its role in microneme secretion during erythrocyte invasion. J Biol Chem. 2013;288(3):1590–602.

9. Kumar S, Kumar M, Ekka R, Dvorin JD, Paul AS, Madugundu AK, et al. PfCDPK1 mediated signaling in erythrocytic stages of Plasmodium falciparum. Nat Commun. 2017;8(1):63.

10. Green JL, Rees-Channer RR, Howell SA, Martin SR, Knuepfer E, Taylor HM, et al. The motor complex of Plasmodium falciparum: phosphorylation by a calcium-dependent protein kinase. J Biol Chem. 2008;283(45):30980–9.

11. Kato N, Sakata T, Breton G, Le Roch KG, Nagle A, Andersen C, et al. Gene expression signatures and small-molecule compounds link a protein kinase to Plasmodium falciparum motility. Nat Chem Biol. 2008;4(6):347–56.

12. Azevedo MF, Sanders PR, Krejany E, Nie CQ, Fu P, Bach LA, et al. Inhibition of Plasmodium falciparum CDPK1 by conditional expression of its J-domain demonstrates a key role in schizont development. Biochem J. 2013;452(3):433–41.

13. Bansal A, Molina-Cruz A, Brzostowski J, Liu P, Luo Y, Gunalan K, et al. PfCDPK1 is critical for malaria parasite gametogenesis and mosquito infection. Proc Natl Acad Sci U S A. 2018;115(4):774–9.

14. Bansal A, Molina-Cruz A, Brzostowski J, Mu J, Miller LH. Plasmodium falciparum Calcium-Dependent Protein Kinase 2 Is Critical for Male Gametocyte Exflagellation but Not Essential for Asexual Proliferation. mBio. 2017;8(5).

15. Kato K, Sudo A, Kobayashi K, Sugi T, Tohya Y, Akashi H. Characterization of Plasmodium falciparum calcium-dependent protein kinase 4. Parasitol Int. 2009;58(4):394–400.

16. Ranjan R, Ahmed A, Gourinath S, Sharma P. Dissection of mechanisms involved in the regulation of Plasmodium falciparum calcium-dependent protein kinase 4. J Biol Chem. 2009;284(22):15267–76.

17. Vidadala RS, Ojo KK, Johnson SM, Zhang Z, Leonard SE, Mitra A, et al. Development of potent and selective Plasmodium falciparum calcium-dependent protein kinase 4 (PfCDPK4) inhibitors that block the transmission of malaria to mosquitoes. Eur J Med Chem. 2014;74:562–73.

18. Kumar S, Haile MT, Hoopmann MR, Tran LT, Michaels SA, Morrone SR, et al. Plasmodium falciparum Calcium-Dependent Protein Kinase 4 is Critical for Male Gametogenesis and Transmission to the Mosquito Vector. mBio. 2021;12(6):e0257521.

19. Dvorin JD, Martyn DC, Patel SD, Grimley JS, Collins CR, Hopp CS, et al. A plant-like kinase in Plasmodium falciparum regulates parasite egress from erythrocytes. Science. 2010;328(5980):910–2.

20. Absalon S, Blomqvist K, Rudlaff RM, DeLano TJ, Pollastri MP, Dvorin JD. Calcium-Dependent Protein Kinase 5 Is Required for Release of Egress-Specific Organelles in Plasmodium falciparum. mBio. 2018;9(1).

21. Kumar P, Tripathi A, Ranjan R, Halbert J, Gilberger T, Doerig C, et al. Regulation of Plasmodium falciparum development by calcium-dependent protein kinase 7 (PfCDPK7). J Biol Chem. 2014;289(29):20386–95.

22. Harper JF, Harmon A. Plants, symbiosis and parasites: a calcium signalling connection. Nat Rev Mol Cell Biol. 2005;6(7):555–66.

23. Abraham EG, Jacobs-Lorena M. Mosquito midgut barriers to malaria parasite development. Insect Biochem Mol Biol. 2004;34(7):667–71.

24. Ishino T, Orito Y, Chinzei Y, Yuda M. A calcium-dependent protein kinase regulates Plasmodium ookinete access to the midgut epithelial cell. Mol Microbiol. 2006;59(4):1175–84.

25. Vontas J, Blass C, Koutsos AC, David JP, Kafatos FC, Louis C, et al. Gene expression in insecticide resistant and susceptible Anopheles gambiae strains constitutively or after insecticide exposure. Insect Mol Biol. 2005;14(5):509–21.

26. Siden-Kiamos I, Ecker A, Nyback S, Louis C, Sinden RE, Billker O. Plasmodium berghei calcium-dependent protein kinase 3 is required for ookinete gliding motility and mosquito midgut invasion. Mol Microbiol. 2006;60(6):1355–63.

27. Davis MI, Patrick SL, Blanding WM, Dwivedi V, Suryadi J, Golden JE, et al. Identification of Novel Plasmodium falciparum Hexokinase Inhibitors with Antiparasitic Activity. Antimicrob Agents Chemother. 2016;60(10):6023–33.

28. Allen JJ, Li M, Brinkworth CS, Paulson JL, Wang D, Hubner A, et al. A semisynthetic epitope for kinase substrates. Nat Methods. 2007;4(6):511–6.

29. Li JL, Baker DA, Cox LS. Sexual stage-specific expression of a third calcium-dependent protein kinase from Plasmodium falciparum. Biochim Biophys Acta. 2000;1491(1-3):341–9.

30. Toenhake CG, Fraschka SA, Vijayabaskar MS, Westhead DR, van Heeringen SJ, Bartfai R. Chromatin Accessibility-Based Characterization of the Gene Regulatory Network Underlying Plasmodium falciparum Blood-Stage Development. Cell Host Microbe. 2018;23(4):557–69 e9.

31. Josling GA, Petter M, Oehring SC, Gupta AP, Dietz O, Wilson DW, et al. A Plasmodium Falciparum Bromodomain Protein Regulates Invasion Gene Expression. Cell Host Microbe. 2015;17(6):741–51.

32. Otto TD, Wilinski D, Assefa S, Keane TM, Sarry LR, Bohme U, et al. New insights into the blood-stage transcriptome of Plasmodium falciparum using RNA-Seq. Mol Microbiol. 2010;76(1):12–24.

33. Trager W, Jensen JB. Human malaria parasites in continuous culture. Science. 1976;193(4254):673–5.

34. Fidock DA, Wellems TE. Transformation with human dihydrofolate reductase renders malaria parasites insensitive to WR99210 but does not affect the intrinsic activity of proguanil. Proc Natl Acad Sci U S A. 1997;94(20):10931–6.

35. Bansal A, Sharma M, Choudhury H. Generation of a new DiCre expressing parasite strain for functional characterization of Plasmodium falciparum genes in blood stages. Sci Rep. 2024;14(1):24076.

36. Chaudhuri S, Seal A, Gupta MD. Autophosphorylation-dependent activation of a calcium-dependent protein kinase from groundnut. Plant Physiol. 1999;120(3):859–66.

37. Oh MH, Wu X, Kim HS, Harper JF, Zielinski RE, Clouse SD, et al. CDPKs are dual-specificity protein kinases and tyrosine autophosphorylation attenuates kinase activity. FEBS Lett. 2012;586(23):4070–5.

38. Kilburn R, Gerdis SA, She YM, Snedden WA, Plaxton WC. Autophosphorylation Inhibits RcCDPK1, a Dual-Specificity Kinase that Phosphorylates Bacterial-Type Phosphoenolpyruvate Carboxylase in Castor Oil Seeds. Plant Cell Physiol. 2022;63(5):683–98.

39. Ahmed A, Gaadhe K, Sharma GP, Kumar N, Neculai M, Hui R, et al. Novel insights into the regulation of malarial calcium-dependent protein kinase 1. FASEB J. 2012;26(8):3212–21.

40. Ito T, Nakata M, Fukazawa J, Ishida S, Takahashi Y. Alteration of substrate specificity: the variable N-terminal domain of tobacco Ca(2+)-dependent protein kinase is important for substrate recognition. Plant Cell. 2010;22(5):1592–604.

41. Asai S, Ichikawa T, Nomura H, Kobayashi M, Kamiyoshihara Y, Mori H, et al. The variable domain of a plant calcium-dependent protein kinase (CDPK) confers subcellular localization and substrate recognition for NADPH oxidase. J Biol Chem. 2013;288(20):14332–40.

42. Harper JF, Breton G, Harmon A. Decoding Ca(2+) signals through plant protein kinases. Annu Rev Plant Biol. 2004;55:263–88.

43. Bansal A, Ojo KK, Mu J, Maly DJ, Van Voorhis WC, Miller LH. Reduced Activity of Mutant Calcium-Dependent Protein Kinase 1 Is Compensated in Plasmodium falciparum through the Action of Protein Kinase G. mBio. 2016;7(6).

44. Andresen C, Niklasson M, Cassman Eklof S, Wallner B, Lundstrom P. Biophysical characterization of the calmodulin-like domain of Plasmodium falciparum calcium dependent protein kinase 3. PLoS One. 2017;12(7):e0181721.

45. Klimecka M, Muszynska G. Structure and functions of plant calcium-dependent protein kinases. Acta Biochim Pol. 2007;54(2):219–33.

46. Solyakov L, Halbert J, Alam MM, Semblat JP, Dorin-Semblat D, Reininger L, et al. Global kinomic and phospho-proteomic analyses of the human malaria parasite Plasmodium falciparum. Nat Commun. 2011;2:565.

47. Fang H, Gomes AR, Klages N, Pino P, Maco B, Walker EM, et al. Epistasis studies reveal redundancy among calcium-dependent protein kinases in motility and invasion of malaria parasites. Nat Commun. 2018;9(1):4248.

